# Modeling cellular response in large-scale radiogenomic databases to advance precision radiotherapy

**DOI:** 10.1101/449793

**Authors:** Venkata SK. Manem, Meghan Lambie, Petr Smirnov, Victor Kofia, Mark Freeman, Marianne Koritzinsky, Mohamed E. Abazeed, Benjamin Haibe-Kains, Scott V. Bratman

## Abstract

Radiotherapy is integral to the care of a majority of cancer patients. Despite differences in tumor responses to radiation (radioresponse), dose prescriptions are not currently tailored to individual patients. Recent large-scale cancer cell line databases hold the promise of unravelling the complex molecular arrangements underlying cellular response to radiation, which is critical to novel predictive biomarker discovery. Here, we present RadioGx, a computational platform for integrative analyses of radioresponse using radiogenomic databases. We first used RadioGx to investigate the robustness of radioresponse assays and indicators. We then combined radioresponse and genome-wide molecular data with established radiobiological models to predict molecular pathways that are relevant for individual tissue types and conditions. We also applied RadioGx to pharmacogenomic data to identify several classes of drugs whose effects correlate with radioresponse. RadioGx provides a unique computational toolbox to advance preclinical research for radiation oncology and precision medicine.

## INTRODUCTION

Radiotherapy is routinely used as curative therapy for cancer patients. Recent technological advances have considerably augmented the physical precision of radiotherapy, resulting in improved cure rates and less toxicity (Baumann et al., 2016; Bernier et al., 2004; Verellen et al., 2007). Biologically motivated improvements (such as the addition of radiosensitizing drugs) to radiotherapy delivery have not seen such dramatic improvements despite the known differences in radiation efficacy that exist among patients with a particular tumor type (Bentzen and Overgaard, 1994; Kozin et al., 2008; Krause et al., 2009). This is due in part to a lack of predictive biomarkers on which to stratify patients. Instead, the stratification of patients to different radiotherapy-containing regimens continues to be based primarily on clinical variables such as tumor stage and patient age (Baumann et al., 2016; Verellen et al., 2007).

The biological determinants of cellular response to radiation, referred to as radioresponse, are complex and include both genomically based cell-intrinsic and external microenvironmental factors (Baumann et al., 2016; Bernier et al., 2004; Steel et al., 1989; Verellen et al., 2007; West et al., 1993). Intrinsic radiosensitivity is thought to vary among individual tumors of the same type with implications for optimal radiotherapy dosing and curability. Measurement of intrinsic radiosensitivity in molecularly-characterized cancer cell lines could provide the radiogenomic data necessary to develop predictors of radioresponse. However, despite decades of research there remain no clinically utilized radiosensitivity biomarkers that have been discovered from cell culture radiogenomic studies. There are many reasons for this, including the need for clonogenic assays when measuring intrinsic radiosensitivity *in vitro* (Puck and Marcus, 1956), which are cumbersome and are not amenable to large screens or radiogenomic studies (Bristow et al., 2018; Yard et al., 2015). Furthermore, radiosensitivity varies with dose in a complex and highly individual manner, rendering measurements at multiple dose-levels a necessity.

Most short-term cytotoxicity assays amenable for high-throughput analysis of drug response have endpoints at 72 hours. These assays are inappropriate for measuring radiosensitivity because of the delayed cellular death by mitotic catastrophe that often occurs following ionizing radiation exposure (Brown and Wouters, 1999). To address this limitation, an extended duration (9-day) viability assay was developed as a surrogate for clonogenic survival that is amenable to high-throughput processing in microtiter plates (Abazeed et al., 2013). This assay was recently applied to 533 cancer cell lines across 17 histologies with multiple radiation dose levels (Yard et al., 2016), becoming the largest radioresponse dataset published by a significant margin. This increase in the scale of radioresponse data holds great potential to contribute to the discovery of robust predictive biomarkers that could someday be translated into clinical use. However, full utilization by the research community requires sophisticated analysis tools that can appropriately model cellular response to radiation and seamlessly integrate associated molecular and pharmacogenomic profiles of cell lines.

In this study, we performed a preclinical assessment of intrinsic radiosensitivity using large-scale radiogenomic datasets (Figure 1). We sought to (*i*) model dose-response data using the linear-quadratic (LQ) model (Brenner, 2008; Dale, 1985; Fowler, 1989); (*ii*) integrate the modeled radioresponse profiles with transcriptomic data to determine pathway- and tissue-specific determinants of radioresponse; (*iii*) infer radioresponse under hypoxic conditions; and (*iv*) identify classes of drugs with cytotoxic effects that correlate with radioresponse. To facilitate these and other future analyses, we developed RadioGx, a new computational toolbox enabling comparative and integrative analysis of radiogenomic datasets. Our work provides a framework for future hypothesis generation and preclinical assessments of radioresponse using appropriate biological assays and indicators.

## RESULTS

### The RadioGx Platform

To realize the full potential of large-scale radiogenomics datasets for robust biomarker discovery, we developed the RadioGx software package (Supplementary Figure 1). RadioGx represents the first computational toolbox that integrates radioresponse data with radiobiologic modeling and molecular data from hundreds of cancer cell lines. Within RadioGx, datasets are standardized with comprehensive cell line annotations including the type of radioresponse assay (i.e., clonogenic assay and 9-day viability assay) and indicators used to generate dose-response data (i.e., SF2 and AUC). RadioGx enables fitting of dose-response data using established radiobiological models, quality control in order to investigate the consistency and biological plausibility of radioresponse assays and indicators, and integration of these data with other data types and radioresponse models (Figure 1).

**Figure 1.**
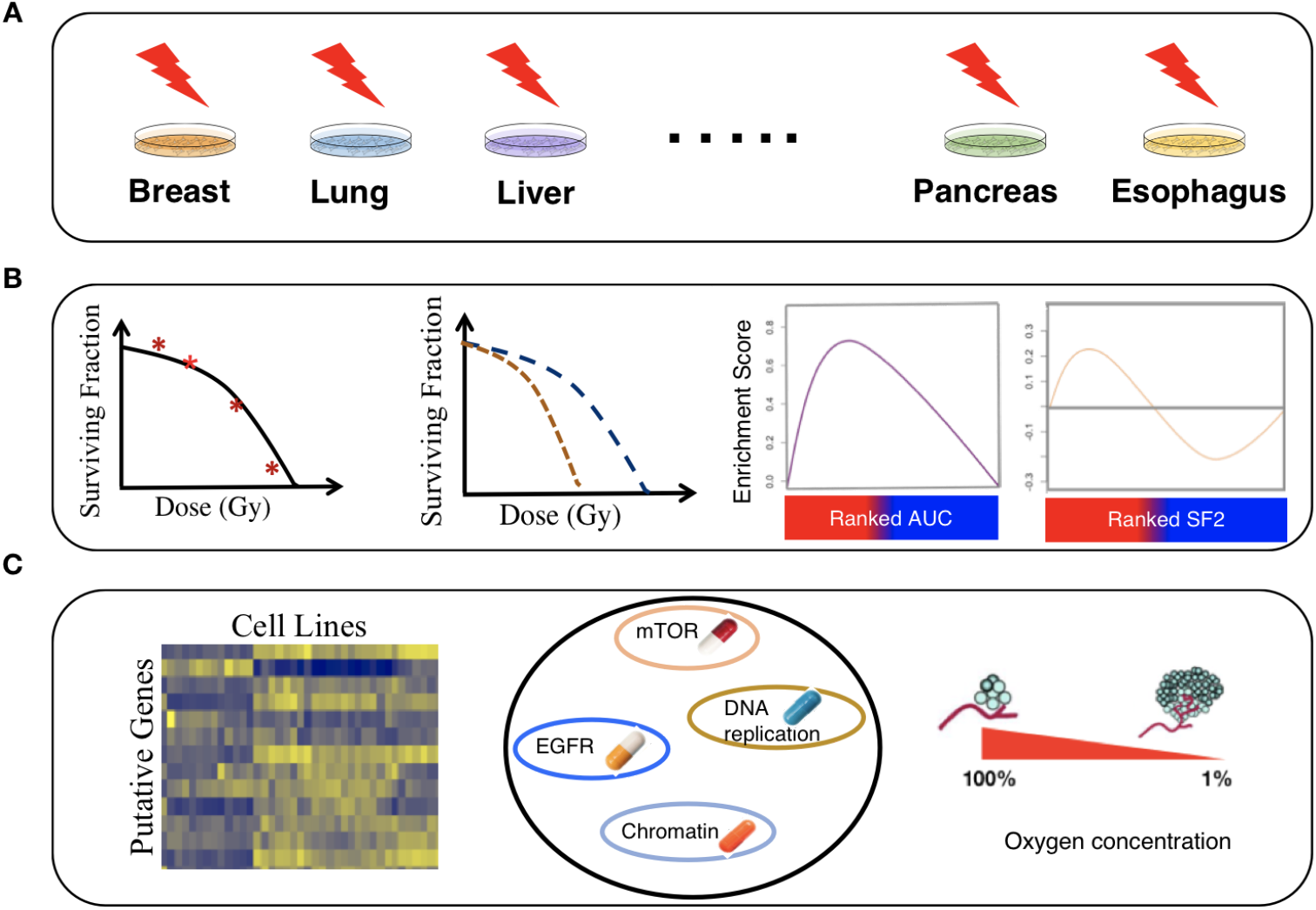
An overview of pre-clinical assessment of radiosensitivity using dose-response and molecular profiling data. (**A**) A schematic illustrating data collection, curation, processing of radiogenomic data obtained from cancer cell lines of diverse histologies. (**B**) A schematic illustrating the process of radiobiological modeling using the linear-quadratic model, assessing the consistency of dose-response data across assays, and evaluating distinct radiation sensitivity indicators such as area under the survival curve (AUC) and surviving fraction after 2 Gy (SF2). (**C**) A schematic illustrating integrative analysis of modeled radioresponse from distinct cell lines with associated genomic/transcriptomic data, pharmacogenomic data, and hypoxia modeling.

### Modeling radiation response within RadioGx

Multiple dose-response measurements from the same cell line can be incorporated into established radiobiological models to predict the effect of specific perturbations (e.g., radiotherapy fraction size or hypoxia) on radioresponse. Within RadioGx, we applied the commonly used linear-quadratic (LQ) model to fit 9-day viability assay data for 533 cancer cell lines (Yard et al., 2016) (Figure 2A). The LQ model goodness-of-fit was high for the majority of cell lines (median R^2^ = 0.958; Supplementary Figure 2A). For 498/533 (93%) of cell lines, the model fit the data reasonably well (R^2^ ≥ 0.6); these cell lines were retained for subsequent analyses.

**Figure 2.**
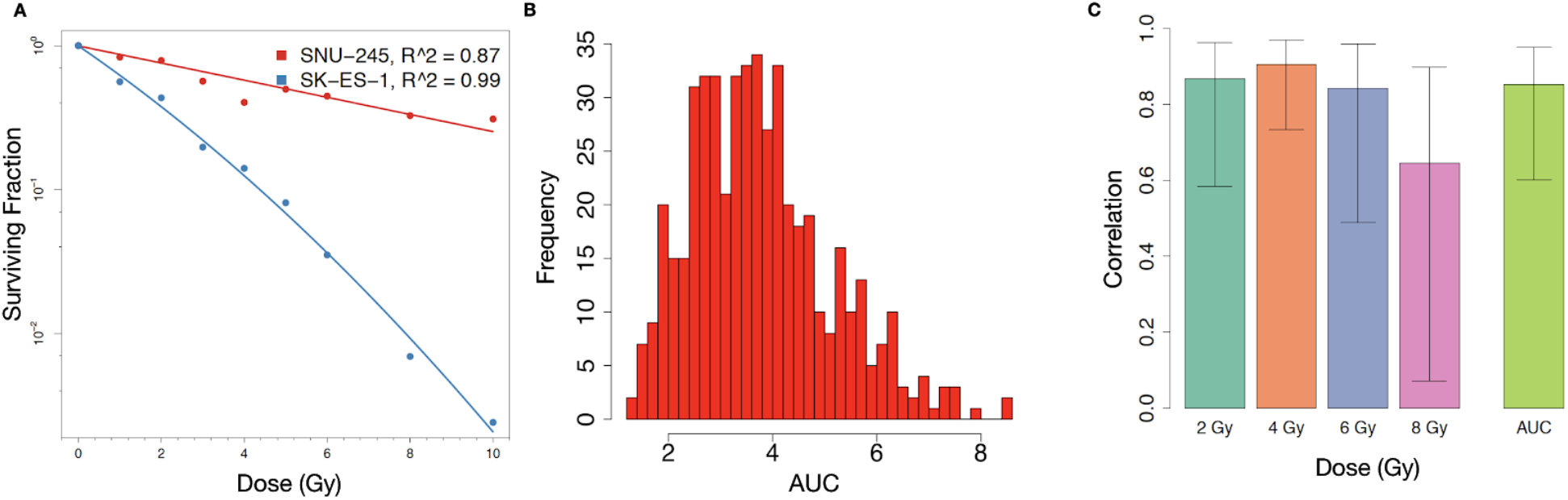
Fitting of dose-response data to the LQ model and concordance of radiation response across asays. (**A**) LQ model fit using RadioGx on the SNU-245 cholangiocarcinoma cell line (red) and SK-ES-1 Ewing sarcoma cell line (blue). The LQ model describes the fraction of cells predicted to survive (y-axis) a uniform radiation dose (x-axis) and is characterized by α and β components for each cell line. For SNU-245 and SK-ES-1, α = 0.14 (*Gy*^-1^),β (*Gy*^-2^) = 0 and α= 0.45 (*Gy*^-1^),β = 0.02 (*Gy*^-2^), respectively. Solid curves indicate the model fit and points denote experimental data (Yard et al., 2016). (**B**) Histogram of AUC values calculated using the *computeAUC* function in RadioGx. (**C**) Correlation (Pearson R with standard deviation) of radioresponse results produced by the 9-day viability assay and the standard clonogenic assay according to the following indicators: SF2, SF4, SF6, SF8, and AUC. Primary data were obtained from Yard et al..

Using the LQ model for each cell line, we calculated the area under the survival curve (AUC) as a summary radioresponse indicator that is independent of a specific dose level. As expected, a range of radioresponse profiles were seen (Figures 2B). We next compared AUC and dose-specific survival data (SF2, SF4, SF6, and SF8) from the 9-day viability assay with clonogenic survival data generated by Yard et al. for a subset of cell lines (Figure 2C and Supplementary Figure 2B-F). We observed high Pearson correlation (R ≥ 0.8) for AUC (n = 15), SF2 (n = 12), SF4 (n = 15), and SF6 (n = 15), but SF8 showed only moderate correlation (R = 0.64; n = 11), consistent with prior observations suggesting poor reproducibility of survival assays following high doses of ionizing radiation (Nuryadi et al., 2018). Taken together, the 9-day viability assay provides a robust surrogate for clonogenic survival to ionizing radiation at a range of dose levels. Moreover, the LQ model within RadioGx allows for characterization of radioresponse and derivation of radioresponse indicators for the vast majority of cancer cell lines.

### Comparison of radioresponse indicators

Summary indicators of radioresponse are useful for preclinical investigations. As radioresponse data within RadioGx has been fit to the LQ model, there is an opportunity to describe radioresponse through imputed survival across a range of dose levels (AUC) or at a specific dose level (e.g., SF2). There is currently no consensus regarding the optimal indicator for use across studies, with both AUC and SF2 frequently used (Deacon et al., 1984; de Jong et al., 2015; Hall et al., 2014; Torres-Roca et al., 2005). The use of SF2 as a radioresponse indicator has been bolstered by clinical observations that local tumor control following radiotherapy may be associated with SF2 measured from *ex vivo* tumor cells (Torres-Roca and Stevens, 2008). Moreover, SF2 is thought to differentiate between radiosensitive and radioresistant cell types (Fertil and Malaise, 1985). However, there is currently insufficient evidence to support the routine use of SF2 or AUC when probing the molecular determinants of radioresponse.

We compared AUC and SF2 across all cell lines within RadioGx. The values were well correlated (ρ = 0.92, 95%CI: 0.90 - 0.93, p=2.2e-16; Figure 3A); the strongest correlations were observed among the most radiosensitive cell lines, and the weakest correlations were observed among the most radioresistant cell lines, where cell death at higher doses likely contributes to the AUC value but has no bearing on SF2 (Figure 3B and Supplementary Figure 3). We then asked whether the biological processes that govern these two radioresponse indicators are the same. To achieve this, we correlated the basal level gene expression data from the Cancer Cell Line Encyclopedia (CCLE) (Barretina et al., 2012) with the radioresponse indicators (SF2 and AUC), and performed gene set enrichment analysis (GSEA) on the gene list ranked based on correlation estimates. For an FDR < 5%, 77 transcriptional pathways were enriched using AUC as the radioresponse indicator, out of which 41 and 36 pathways were positively and negatively correlated with AUC, respectively (Supplementary File 1, Supplementary Figure 4). Similarly, using SF2 as the radioresponse indicator, only 38 pathways were enriched, out of which 19 were positively correlated with the SF2 value. All but three of the pathways enriched using SF2 were enriched using AUC (Figures 3C and 3D).

The 17 pathways that were significantly correlated with radioresponse using the AUC indicator but not the SF2 indicator included biological processes known to impact radioresponse, suggesting stronger relevance for AUC. For instance, the *Nrf2-mediated oxidative stress response* pathway was positively associated with AUC but not with SF2 (Supplementary File 1). In conditions of oxidative stress, such as following radiation, degradation of Nrf2 is prevented, leading to its stabilization and translocation into the nucleus, where it activates expression of a wide variety of downstream antioxidant targets (Espinosa-Diez et al., 2015); this pathway has previously been described as contributing to intrinsic radioresistance (Abazeed et al., 2013; Singh et al., 2010). In addition, progression through the cell cycle following radiation response is a known factor in determining cell survival vs. cell death via mitotic catastrophe. Three pathways directly related to cell cycle progression ([1] *cell cycle: G2/M DNA damage checkpoint regulation*, [2] *cell cycle: G1/S checkpoint regulation*, and [3] *cell cycle control of chromosomal replication*) were all seen exclusively when using AUC as the radioresponse indicator. Thus, as compared with SF2, AUC was able to capture more gene expression pathways putatively correlated with radioresponse including pathways with known mechanistic roles in mediating cellular radioresponse. Taken together, our analyses reveal AUC and SF2 as related radioresponse indicators with AUC providing for a more comprehensive characterization of the biological processes underpinning radioresponse. As a result of these findings, we exclusively used AUC as the radioresponse indicator for subsequent analyses.

**Figure 3.**
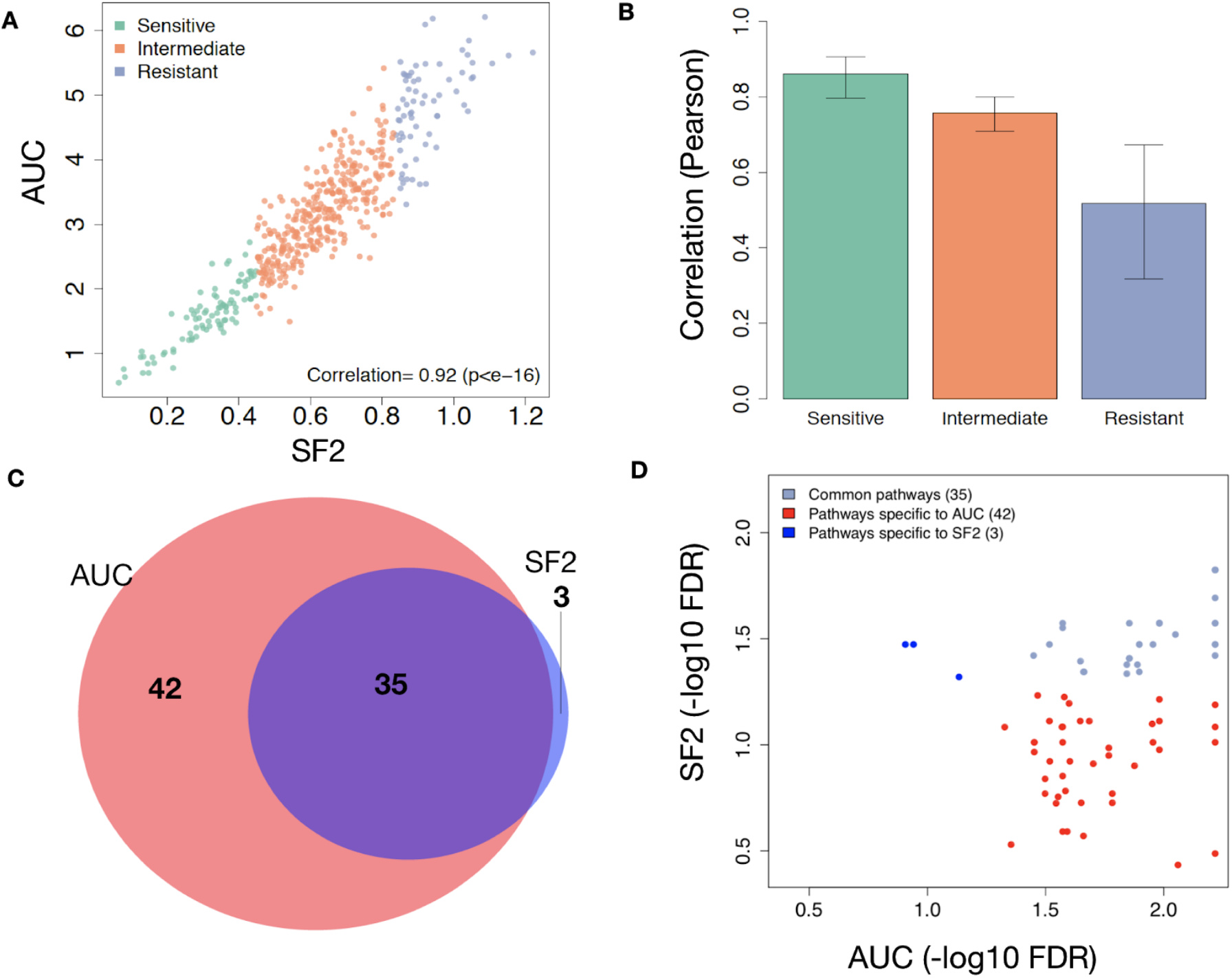
Concordance of SF2 and AUC. **(A)** Correlation between the radioresponse indicators, SF2 and AUC, across 498 cell lines. **(B)** Pearson correlation (with standard deviation) between SF2 and AUC across 498 cell lines based on tertiles. **(C)** Venn diagram illustrating the transcriptional pathways associated with radioresponse using SF2 or AUC as the response indicator. **(D)** False discovery rate (FDR) for each transcriptional pathway from (C) illustrating greater levels of statistical significance among pathways specific to AUC.

### Radiobiological modeling to estimate impact of DNA repair on survival

The LQ model can be used to estimate the dependence of cellular survival on radiotherapy fraction size and DNA repair. The α and β values produced by the LQ model allow for comparisons among distinct cell lines or tumors, and in clinical practice the α/β ratio is used to predict cellular response to different radiotherapy fractionation schemes.

Using the LQ model, we derived the α/β ratio for cancer cell lines within RadioGx. A wide range of α/β values were observed (Figure 4A; median = 10.14; interquartile range = 4.49 - 28.07). As expected, the α component was strongly anti-correlated with AUC, whereas the β component displayed no significant association with AUC (Figure 4B). This result indicates that for the cell line data contained within RadioGx, dependence of cellular survival on radiotherapy fraction size is a distinct parameter that describes radioresponse and should therefore be considered alongside radiosensitivity (e.g., AUC or SF2) in preclinical investigations.

In order to examine the biological factors that underlie the differences between α, β and AUC, we identified transcriptional pathways that were significantly associated with each radioresponse metric. For an FDR of 5%, we found 14 pathways commonly associated with all 3 metrics (Figure 4C; Supplementary File 2). Supporting the biological relevance of these pathways, several known components of DNA damage response, signaling, and repair were represented among the 14 common pathways. For instance, pathways related to *mismatch repair in eukaryotes, role of BRCA1 in DNA damage response*, and *cell cycle control of chromosomal replication* were each present. These results, which are consistent with fundamental tenets of radiobiology, suggest that analysis of large cell line resources within RadioGx could be performed to generate novel hypotheses and could contribute to preclinical biomarker discovery.

**Figure 4.**
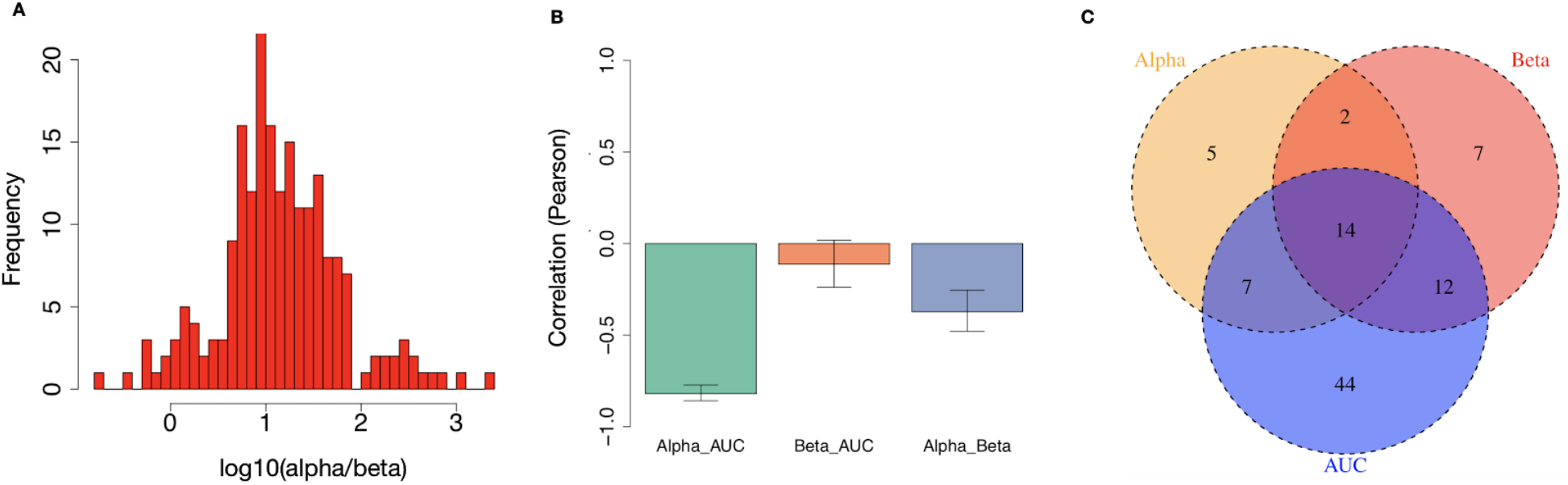
Distinct biological underpinnings of α/β derived from the LQ model. **(A)** Histogram of α/β (*Gy*) values obtained from the LQ model across all cell lines. **(B)** Pearson correlations (with standard deviation) between AUC and the α and β components of the LQ model. **(C)** Transcriptional pathways that are significantly associated with AUC, α, and/or β.

### Modeling the effects of hypoxia on radioresponse

By integrating radioresponse and molecular data, RadioGx is meant to enable new biological insights and predictions. To further demonstrate the utility of RadioGx for this purpose, we next extended the radiobiological modeling to incorporate the putative effects of oxygen availability in the tumor microenvironment on radioresponse (Daşu et al., 2005).

Molecular oxygen is necessary to mediate the indirect effects of ionizing radiation to exert cell kill. Thus, cells become more resistant to radiation under oxygen-deficient conditions. We derived adjusted AUC values for the cancer cell lines within RadioGx at a range of oxygen partial pressures. As expected, reduced oxygen partial pressure resulted in a predicted increase in AUC (Figure 5A). Cell lines from distinct cancer histologies displayed consistent increases in AUC under hypoxic conditions (p<2.2e-16 for all, Wilcoxon test), but the magnitude of this increase differed between histologies (Figure 5B). The largest and smallest median differences in AUC were observed for cancer cell lines from the breast and large intestine, respectively. These differences reflect a non-linear relationship between oxygen availability and radioresponse that is dependent on α/β.

**Figure 5.**
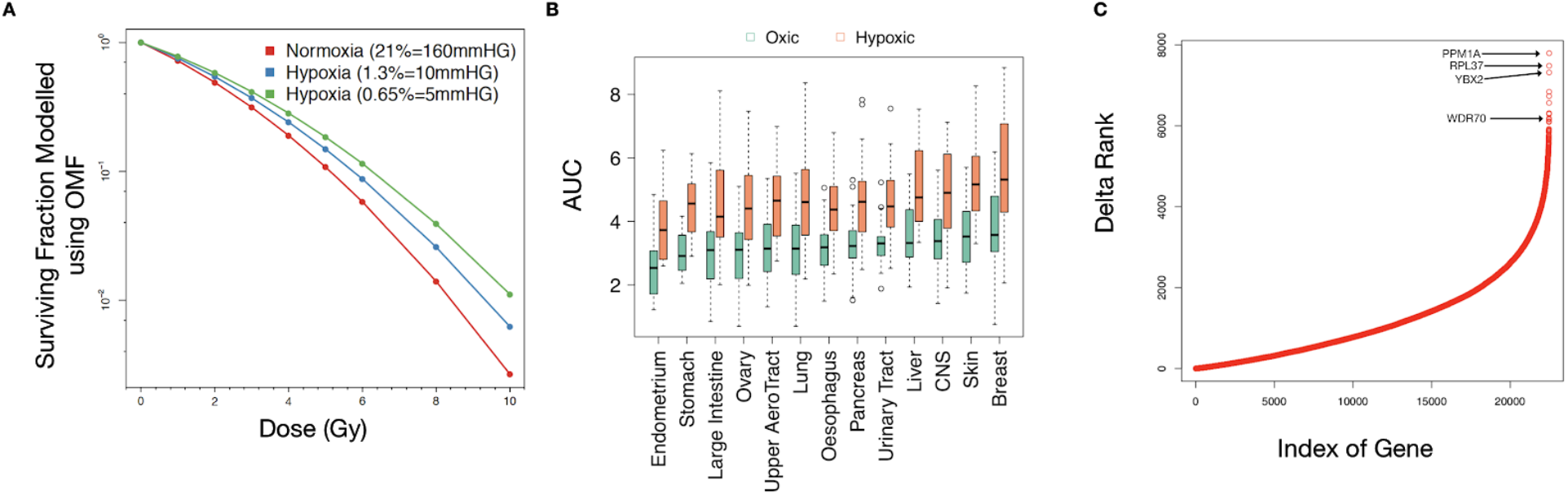
Integrative analysis of radiobiological model with transcriptomic data and prediction of radioresponse under hypoxia. **(A)** Hypothetical illustration of cancer cell surviving fraction according to dose and oxygen partial pressure, as modeled using RadioGx. Solid curves are modeled using Equation (3) (Methods). The computed AUC values are 2.41, 2.71, 2.97 for normoxia (160 mmHg), hypoxic condition with 10 mmHg, and hypoxic condition with 5 mmHg, respectively. **(B)** Changes in AUC by tissue type (with minimum of 15 cell lines within RadioGx) under normoxic (160 mmHg) or hypoxic (5 mmHg) conditions, ordered according to median AUC under normoxia. **(C)** The difference in ranks are shown between the strength of univariate association of each gene with AUC under normoxic (160 mmHg) vs. hypoxic (5 mm Hg) conditions across cancer cell lines within RadioGx.

Next, we evaluated the univariate association of gene expression levels measured under normoxic conditions with AUC values under normoxic and hypoxic conditions. For an FDR < 5%, the numbers of genes that were significantly associated with radioresponse were 1,825 and 2,395 under normoxic and hypoxic conditions, respectively (Supplementary File 3). Moreover, 1,375 genes were negatively associated with radioresponse under normoxic condition but positively associated with radioresponse under hypoxic condition, and 471 genes were positively associated with radioresponse under normoxic condition but negatively associated with radioresponse under hypoxic condition (Supplementary Figure 5). In keeping with these effects, we observed large changes in the ranking of strength of correlation of specific genes with radioresponse under oxic and hypoxic conditions (Figure 5C). The gene with the greatest change, PPM1A, has been implicated in the regulation of cellular stress response and has previously been shown to have hypoxia-specific activity (Heikkinen et al., 2010). WDR70, a gene with known roles in DNA double strand break repair (Guo et al., 2016; Zeng et al., 2016), also displayed a large change in this analysis (Figure 5C). One might hypothesize based on our results that WDR70 could have previously uncharacterized hypoxia-specific activities and/or expression; these findings warrant further investigation.

### Tissue specificity of radioresponse and repair

It is known that distinct tissues and tumor types respond differently to ionizing radiation exposure. Intrinsic radiosensitivity has been suggested as a major contributing factor to this differential response (Yard et al., 2016). We used RadioGx to interrogate radioresponse within tissue types represented by a minimum of 15 cell lines (Figure 6).

To examine the biological factors that may underlie suspected differences in radioresponse between tissue types, we identified 281 transcriptional pathways that were significantly associated with radioresponse within at least one tissue type (Figure 6A; Supplementary File 4). Of these 281 pathways, 123 were statistically significant only in one tissue type (Supplementary Figure 6). Overall, there were more statistically significant pathway associations with radiosensitivity than radioresistance (total across all tissue types: 437 and 226, respectively). Remarkably, we did not find any transcriptional pathways that were statistically significantly associated with radioresponse across all tissue types. We also observed variable α/β values among the tissue types within RadioGx (Figure 6B), suggesting heterogeneity of DNA repair and dependence on radiotherapy fraction size. Our findings support the use of tumor-specific as opposed to pan-cancer radioresponse biomarkers and radiosensitizing strategies.

**Figure 6.**
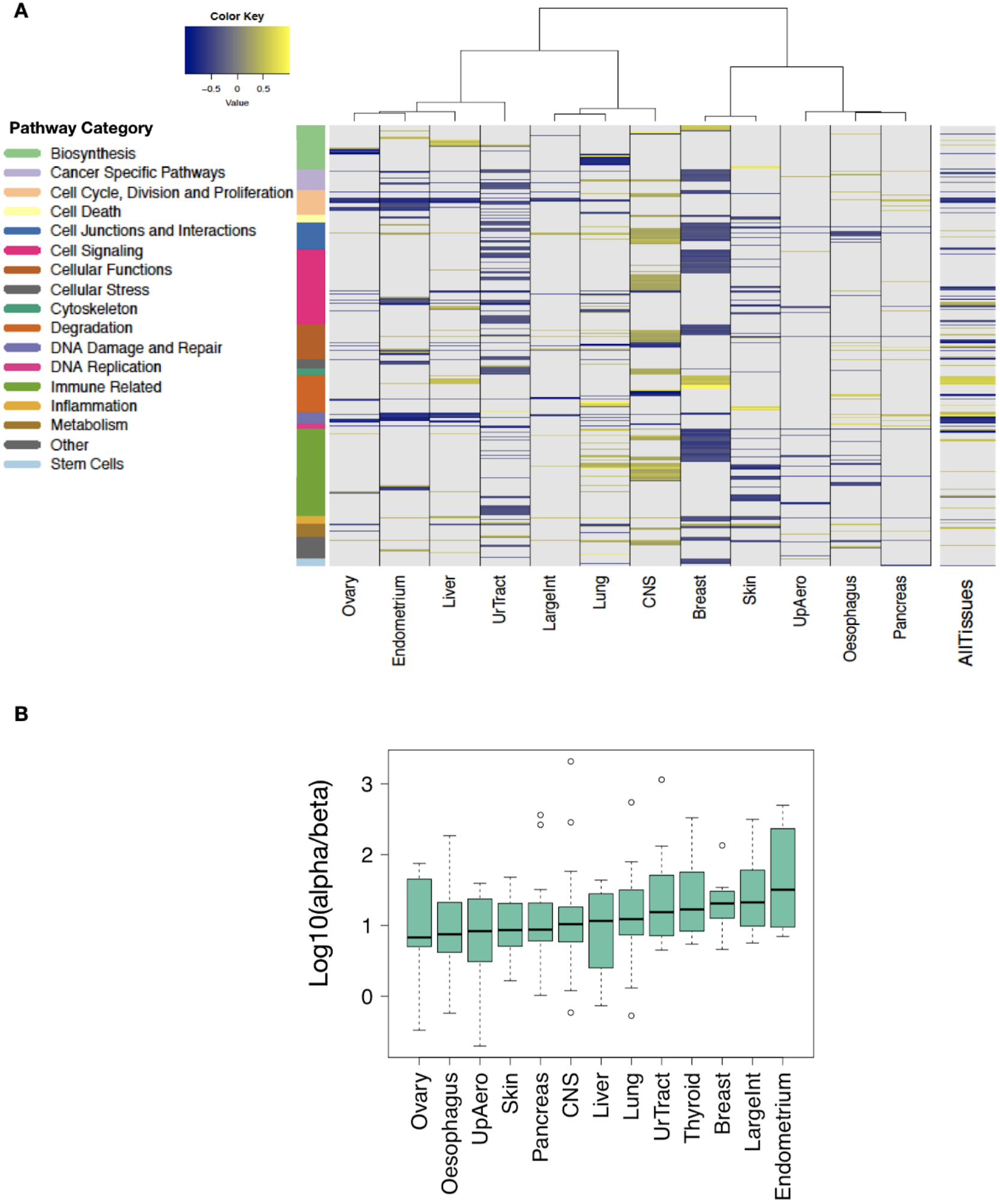
Tissue specificity of molecular determinants of radioresponse. **(A)** The tissue types (columns) represented by a minimum of 15 cancer cell lines were considered for analysis. A total of 281 pathways are depicted (rows) and are annotated by function. Colours designate pathways significantly associated with AUC (FDR < 5%). **(B)** Heterogeneity of α/β ratios across cancer cell lines derived from distinct tissue types ordered according to median values.

### Common dependencies of therapeutic effects among radiotherapy and drugs

Datasets within RadioGx are standardized with regard to cell line annotations such that integrated analyses using other existing datasets can be easily conducted. For instance, our previously published tool, PharmacoGx (Smirnov et al., 2016), contains pharmacogenomic data from multiple studies and enables meta-analysis of pharmacogenomic data. We wished to identify categories of drugs with cytotoxic effects that correlate with radioresponse, so we interrogated RadioGx to compare cellular responses to ionizing radiation and chemotherapeutic agents (n=545 distinct drugs). Drug responses were obtained from 480 cancer cell lines from the CTRPv2 pharmacogenomic dataset (Supplementary Table 1) that were in common between the datasets. We computed the correlation between drug response and radiation response across the cancer cell lines (Supplementary Figure 7) and then classified drugs according to pharmacological categories (i.e., by cellular targets and/or mechanisms of action). Drugs targeting the cytoskeleton, DNA replication, and mitosis displayed the strongest correlations with radioresponse (FDR < 5%) (Figure 7). Thymidylate synthetase inhibitors such as the known radiosensitizing drug, fluorourocil, also displayed cytotoxic effects that correlated with radioresponse but did not reach statistical significance. In addition to these anticipated and largely confirmatory findings, we also observed unexpected negative associations between radioresponse and cytotoxic effects of drugs targeting numerous cell signaling pathways (i.e., PI3K signalling, ERK MAPK signaling, WNT signalling, EGFR signalling, ABL signalling), although these were not statistically significant.

**Figure 7.**
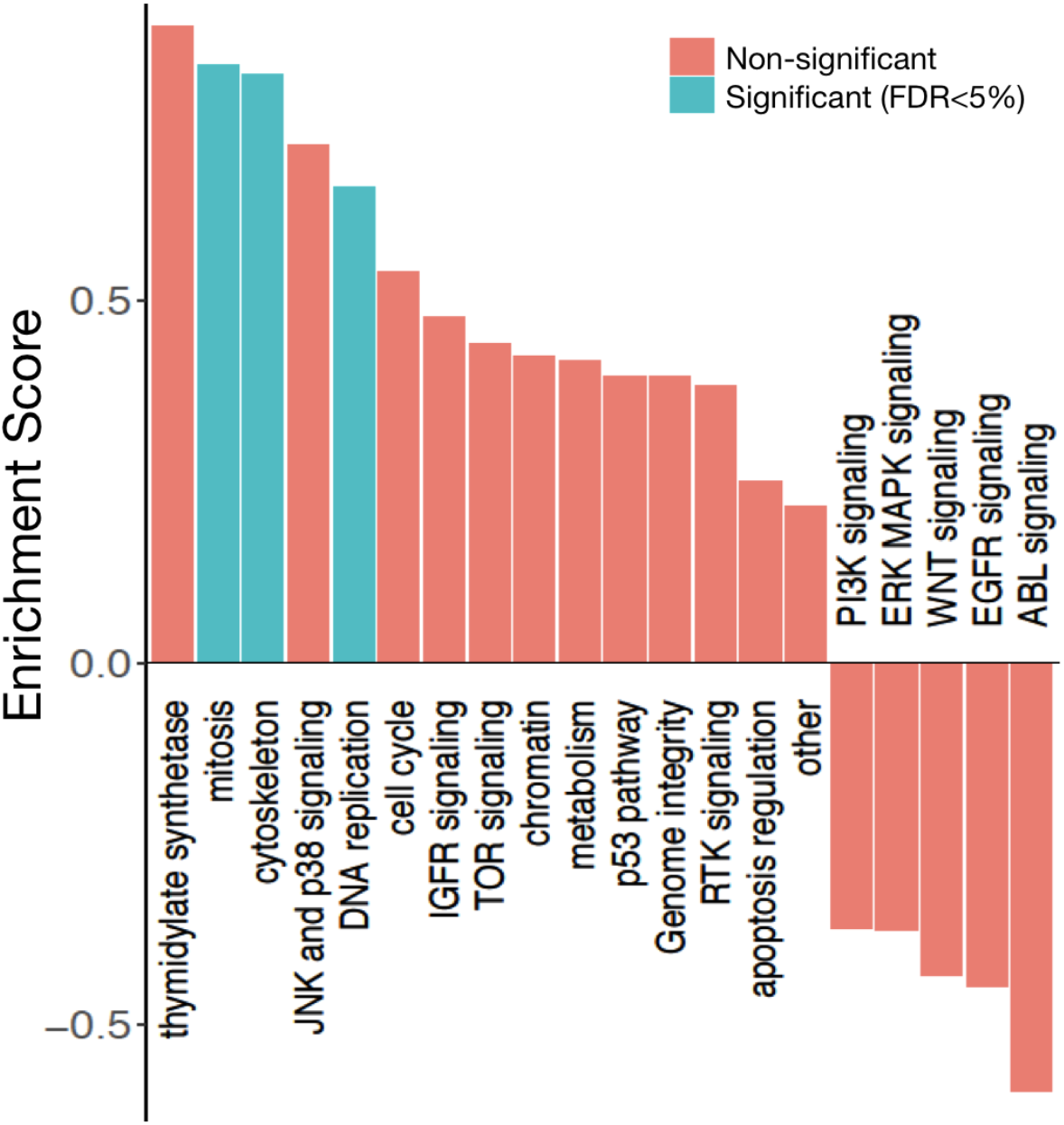
Identification of drugs and pharmacological classes with cytotoxic effects on cancer cell lines that correlate with radioresponse. Pharmacological enrichment analysis using radiation AUC as the radioresponse indicator. Pharmacological classes with statistically significant associations with radioresponse in cancer cell lines are indicated.

## DISCUSSION

To date, the paradigm of precision medicine has primarily been applied to advanced incurable cancers. For early stage curable cancers for which radiotherapy is used with curative intent, there remains a need for more precise biologically-tailored radiotherapy delivery. For instance, there are currently no clinically implemented molecular biomarkers that are predictive of radioresponse. This also extends to predictive insights into the response of tumors to other therapeutic agents that may be administered in combination with radiotherapy. Although molecular diagnostic tools are making their way into clinical practice in other settings, the lack of equivalent molecular indicators in the field of radiobiology has impeded translation in this domain (Baumann et al., 2016; Bibault et al., 2013; Bristow et al., 2018).

Recently, large radioresponse and genomic datasets have been generated from hundreds of cancer cell lines, providing an opportunity to address this unmet need. We have developed RadioGx, an open-source software package that enables users to perform integrative analysis of radiogenomic datasets for preclinical evaluation of radioresponse determinants. RadioGx standardizes published nomenclature and annotations between datasets and integrates dose-response and molecular data.

We used RadioGx to evaluate the appropriateness of the 9-day viability assay for assessing radioresponse, the robustness of distinct radioresponse indicators, and the utility of applying established radiobiological models to the data for novel hypothesis generation. We confirmed the findings from Yard et al. that the 9-day viability assay, which is amenable to high-throughput processing and analysis, largely recapitulates the results of the more tedious clonogenic assay. We note that some prior putative intrinsic radiosensitivity gene expression signatures that were generated using cell line clonogenic survival data have failed to validate using independent sets of cancer cell lines (Bratman et al., 2017; Hall et al., 2014), highlighting the need for robust reproducible methodologies for future studies. Moreover, we found that AUC derived from the LQ model might provide a more complete characterization of the biological processes underpinning radioresponse as compared with the dose-specific SF2 indicator, particularly for relatively radioresistant cell types. Based on our findings, we suggest that AUC be the radioresponse indicator of choice for preclinical studies. While we found that the LQ model fit the radioresponse data for the vast majority of cancer cell lines within RadioGx, a small subset were not amenable to LQ modeling and should be excluded or used with caution in future radiogenomic analyses.

A major hurdle in the development of large-scale radioresponse datasets has been the technical and throughput challenges associated with the clonogenic assay. We demonstrated how existing data within RadioGx can be used to generate hypotheses and make predictions to inform future investigations. For instance, recognizing a dearth of large-scale radioresponse data under hypoxic conditions, we integrated radiobiological modeling with gene expression data from RadioGx, which allowed us to predict radioresponse under hypoxic conditions. Our findings suggest that the change in radioresponse under hypoxia is tissue-specific and that specific genes are either differentially associated with radioresponse under normoxic and hypoxic conditions or may have expression levels or activity that are regulated by oxygen tension; these specific hypotheses generated by our analysis through RadioGx could be tested experimentally in future studies. In addition, by combining RadioGx with an existing pharmacogenomics analysis platform, we uncovered drugs with cytotoxic effects that are correlated or anti-correlated with radioresponse, suggestive of genomic/transcriptomic dependencies related to their mechanisms of action. We were able to confirm drug classes with therapeutic effects that overlap with ionizing radiation (e.g., mitotic inhibitors); moreover, this analysis proposed novel hypotheses regarding possible anticorrelated therapeutic effects with drugs targeting a number of cellular signaling pathways such as ABL and EGFR. Future studies may seek to examine whether members of these drug classes may make rational combination therapies with radiation as a result of reduced additive toxicity.

In summary, this study demonstrates the impact of combining radiogenomic datasets with established radiobiological models and other existing pharmacogenomic data. Future applications of RadioGx may include generation of biomarkers for intrinsic radiosensitivity and selection of novel combination therapies for preclinical testing. Thus, we envision that RadioGx will help to accelerate preclinical radiotherapeutic discovery pipelines and guide the selection of appropriate biological endpoints.

## METHODS

### Curation of dose-response and transcriptomic data

One of the major hurdles in genomic studies involving cell lines is the lack of standardized identifiers for cell lines. In order to overcome this, we assigned a unique identifier to each cell line and radiation therapy, and matched entities with the same unique identifier throughout the implementation. Moreover, there is a lack of standardization in annotating genomics features, i.e. annotating probe expression to gene expression across various microarray datasets. Hence, we have used the annotations from the BrainArray database, which reflect recent annotation of the human genome to perform the mapping from microarray probe sets to genomic expression data.

We implemented a RadioSet (also known as RSet) in the RadioGx package. This class is a data container storing radiation dose-response and molecular data along with experimental metadata (detailed structure provided in the Supplementary Materials). In addition, this class also enables efficient implementation of curated annotations for cell lines, and molecular features, which facilitates comparisons between across different datasets. We have implemented a unique set of functions that facilitates users to analyze radiogenomic datasets. One of the primary functions is the *downloadRSet* that allows users to download the *RadiationSet* (RSet) object. We have also incorporated a function, *linearQuadraticModel*, which plots the radiation cell survival curve using the standard radio-biological formalism, the linear-quadratic (LQ) model (see below). This function considers normal distribution to fit the errors as a default parameter, but the users also have an option to choose Cauchy distribution. For a given dataset by the end user, this function fits the dataset with the LQ model, and returns radiobiological parameters alpha and beta along with the goodness of fit. To extract several features from this curve, we have implemented a function *computeAUC*, which enables users to compute area under the survival curve (AUC), *computeSF2* function, which returns the fraction of cells that survive a radiation dose of 2Gy, *computeD10* function, which returns the radiation dose at which only 10% of cells survive.

Supplementary Table 1 presents the sensitivity and transcriptomic datasets that are used in this study, and the functionality of the RadioGx package is presented in Table 1.

**Table 1.**
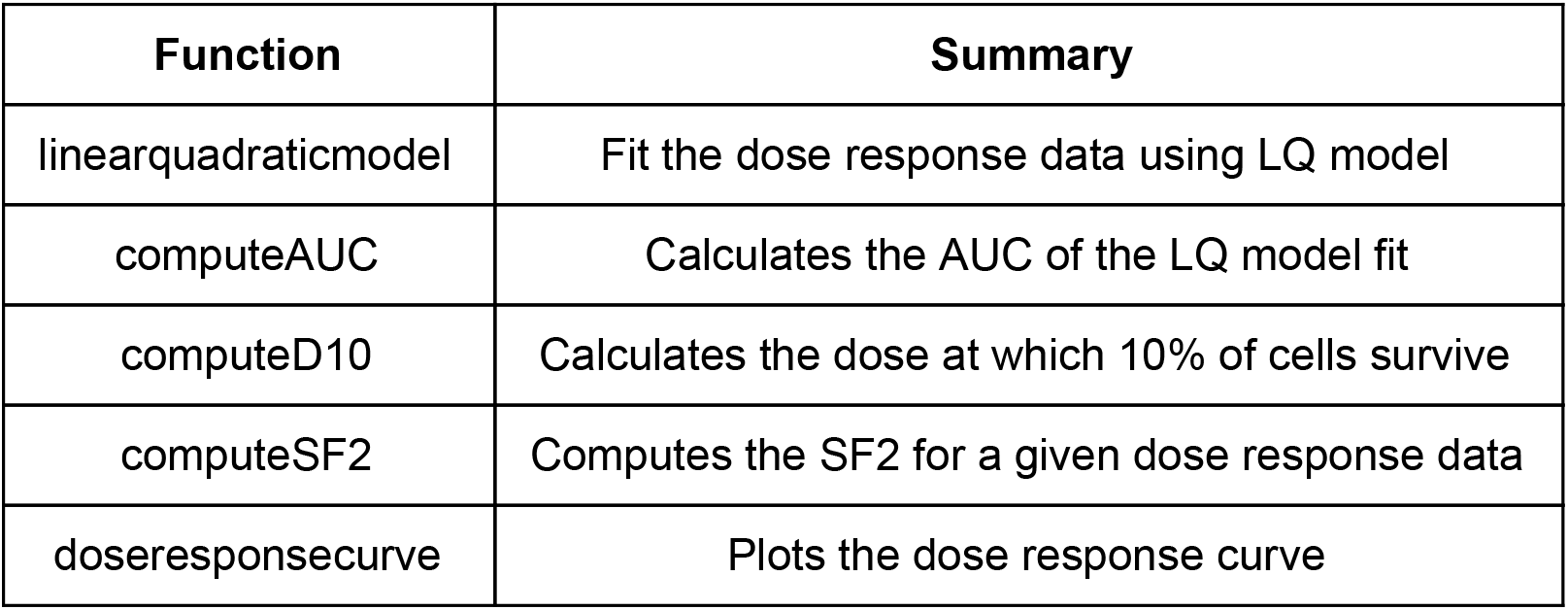
Functionality of RadioGx package.

### Radiobiological Model

A radiobiological model is a formulation that is used to allow comparisons of various clinically relevant radiotherapy treatment regimens. The most commonly used model in current clinical practice is the LQ model (Brenner, 2008; Dale, 1985; Fowler, 1989), which assumes that there are two components to cell killing induced by radiation: one that is proportional to dose (linear, α) and another that is proportional to the square of the dose (quadratic, β). The LQ model describes the fraction of cells that survived (S) a uniform dose D (Gy); the survival fraction of cells after irradiating with an acute dose D is given by:

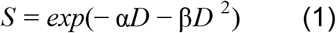

The ratio 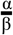 varies by the cell population or tissue that is being irradiated, and reflects the response to different fractionation schemes. Cell populations or tissues with a high value are less sensitive to the effects of fractionation than those with a low value.

### Radiobiological Modelling of Hypoxia

The LQ model can also be used to model the effect of hypoxia. Hypoxia is a hallmark of many solid malignant tumors and influences tumor progression, therapy resistance, development of metastases, clinical behavior, and response to conventional treatments like radiotherapy (Hall and Giaccia, 2012). The survival fraction of cells due to a given radiotherapy dose is given by Equation (1) under well-oxygenated, or normoxic conditions. However, the surviving fraction of cells may vary depending on the amount of oxygen concentration in the tumor, as cells in the hypoxic region are considered to be more resistant to radiation therapy. This hypoxic effect can be incorporated into the LQ model using the, *“Oxygen Enhancement Ratio (OER*)”, which can be normalized to yield the *“Oxygen Modification Factor (OMF*)” (Alper and Howard-Flanders, 1956; Daşu et al., 2005; Titz and Jeraj, 2008; Wouters and Martin Brown, 1997). OMF is defined as follows:

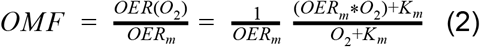

where *O*_2_ is the oxygen concentration in the system in mmHg, *K_m_* = 3 mm Hg, defined as the oxygen at which half of the ratio is achieved, and *OER_m_* = 3 is the maximum value at well-oxygenated condition. Therefore, the LQ model given in Equation (1) can be modified to include oxygen concentration as follows:

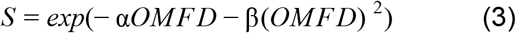

In general, the OER can be a function of radiation dose. Some studies have suggested that the maximal oxygen enhancement varies in the range of 2.5-3 with differences in radiation dosage (Freyer et al., 1991; Palcic and Skarsgard, 1984; Skarsgard and Harrison, 1991). This can be simply included into the revised LQ model by considering different OERs for the parameters α and β, that is, *OER*_α_ and *OER*_β_. However, since we consider the normalized OER (or, OMF), the introduction of these separate terms will not produce a significant difference in the final survival fraction. Thus, we assume *OER*_α_ = *OER*_β_ in our mathematical framework. We assume that the system is moderately hypoxic, i.e. approximately 5 mm HG for the present study.

### Association with drug response and Pharmacological Enrichment Analysis

We used CTRPv2 dataset in PharmacoGx package (version 1.10.3) (Smirnov et al., 2016) that has 545 drugs to compute the association between radioresponse and drug response (defined by the Area under the curve of the Hill function). We also performed pharmacological enrichment analysis, an adaptation of the GSEA methodology. For this, we computed the correlation of radioresponse with each drug response, and a pharmacological set represents a gene set. Similar to the GSEA method, a running sum is calculated, starting with the first compound-level statistic to the last. The sum is increased if a compound-level statistic belongs to the pharmacological class of interest, otherwise, the sum is decreased. The enrichment score of the pharmacological class of interest is defined as the maximum deviation from zero of the running sum (Seashore-Ludlow et al., 2015) (Supplementary Figure 8).

### Pathway Analysis

The pathway enrichment analysis on the gene expression data is carried out using the gene set enrichment analysis (GSEA) method (Subramanian et al., 2005) with pathways defined by QIAGEN’s Ingenuity^®^ Pathway Analysis (IPA^®^, QIAGEN Redwood City, www.qiagen.com/ingenuity). Genes were ranked based on their coefficient of correlation between the gene expressions and the IHC scores (core density or stromal retention ratio). GSEA was then used to compute the enrichment score for each pathway with statistical significance calculated using a permutation test (10,000 permutations) as implemented in the *piano* package (Väremo et al., 2013). Nominal p-values obtained for each pathway are corrected for multiple testing using the false discovery approach (FDR) (Benjamini and Hochberg, 1995).

### Research Reproducibility

RadioGx is implemented in R. The code, documentation, and detailed tutorial describing how to run our pipeline and reproduce our analysis results are open-source and publicly available through the RadioGx GitHub repository (https://github.com/bhklab/RadioGx). A virtual machine reproducing the full software environment is available on Code Ocean. Our study complies with the guidelines outlined in (Gentleman, 2005; Sandve et al., 2013; Stroup et al., 2000). All the data are available in the form of RSet objects with associated digital object identifiers (DOI).

## ACKNOWLEDGMENTS

This work was supported by a grant to SVB from the V Foundation for Cancer Research (V2018-010). SVB and BHK are supported by the Gattuso-Slaight Personalized Cancer Medicine Fund at the Princess Margaret Cancer Centre. ML is supported by a fellowship from STARS21. We also gratefully acknowledge the support from the Princess Margaret Cancer Foundation and the Princess Margaret Cancer Center Head & Neck Translational Program, with philanthropic funds from the Wharton Family, Joe’s Team, and Gordon Tozer.

---

**Supplementary Figure 1.**
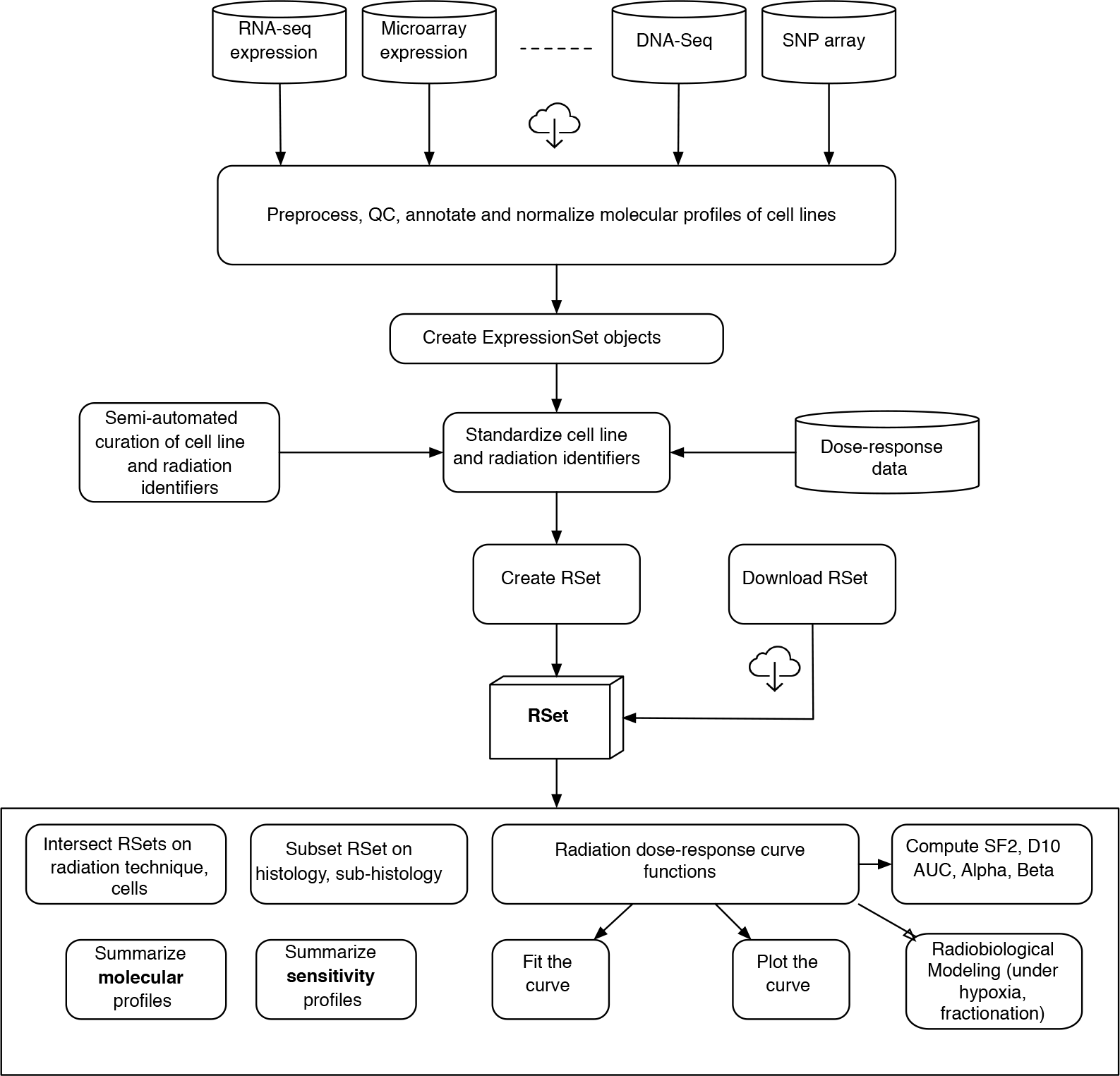
Design of RadioGx platform.

**Supplementary Figure 3.**
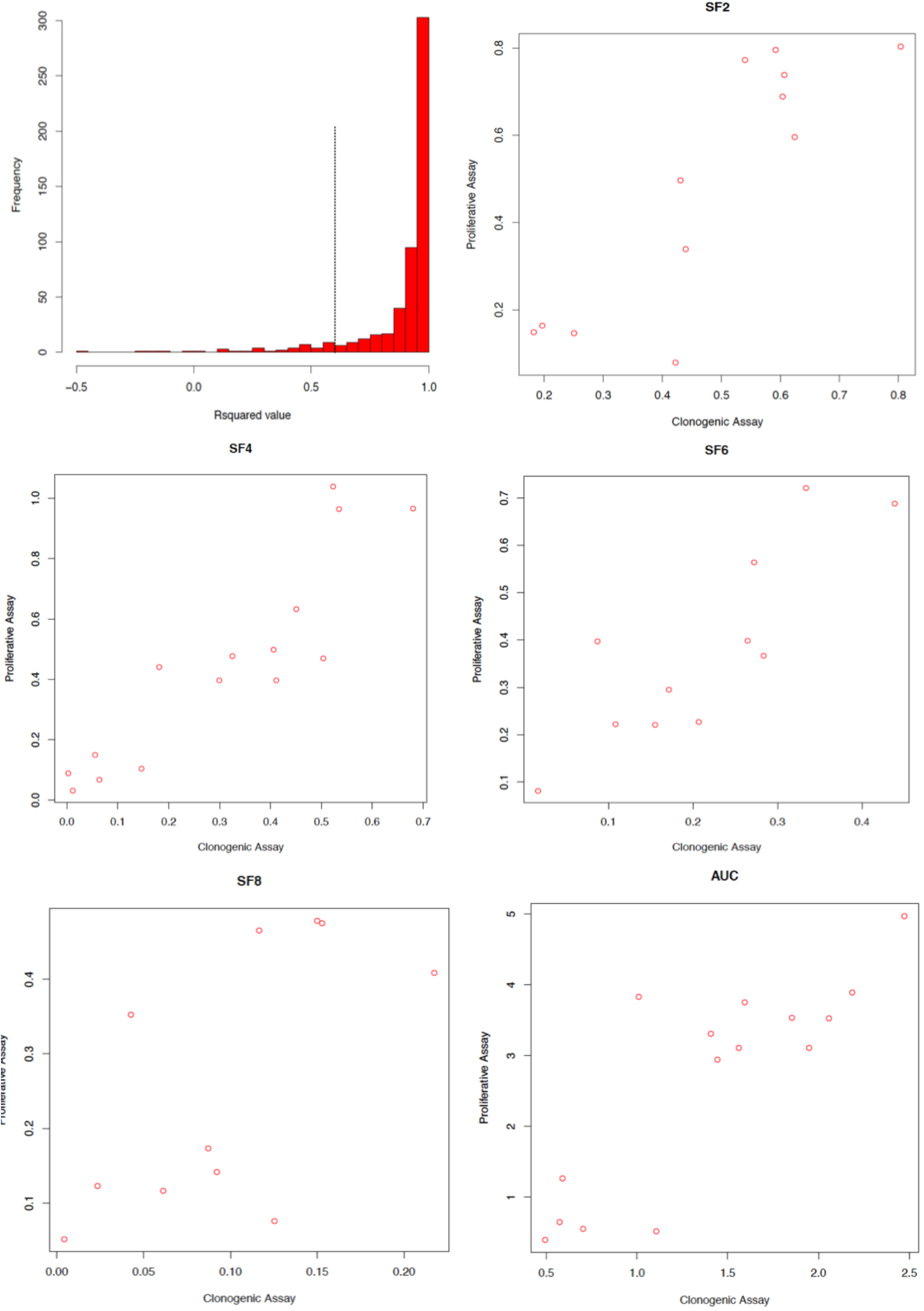
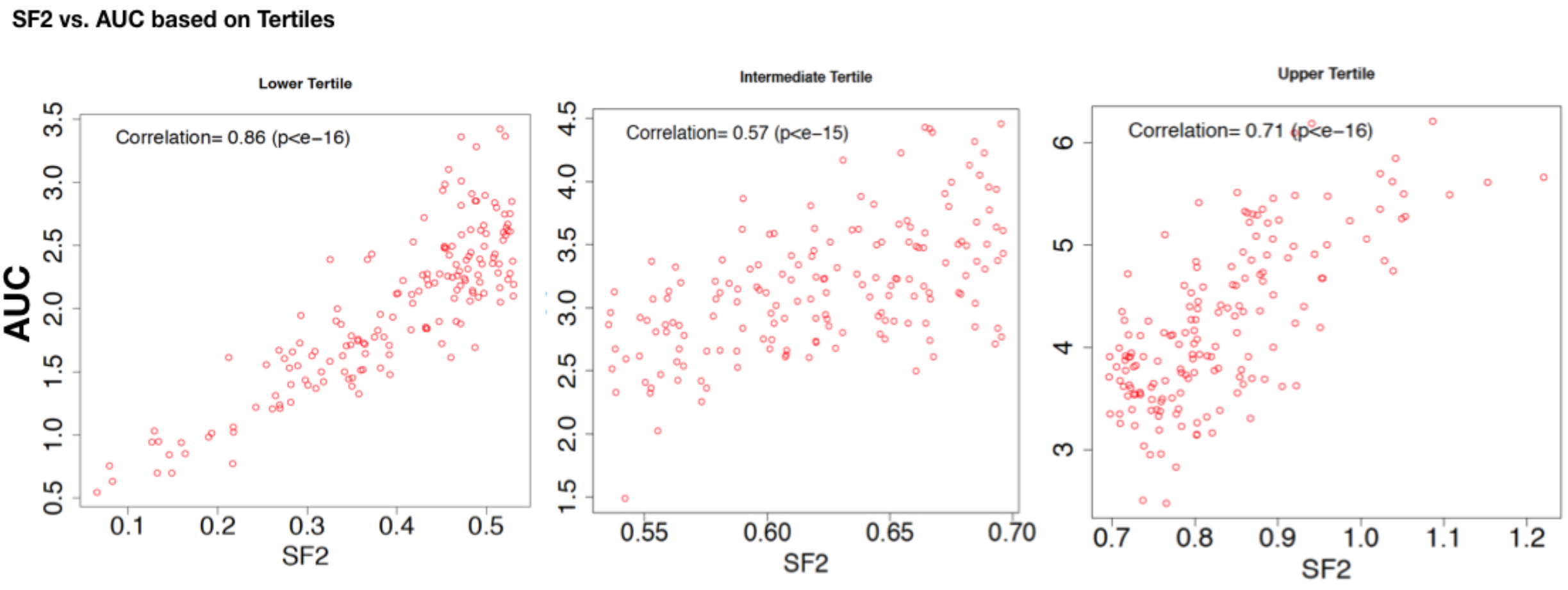
Comparison of SF2 and AUC based on tertiles.

**Supplementary Figure 4.**
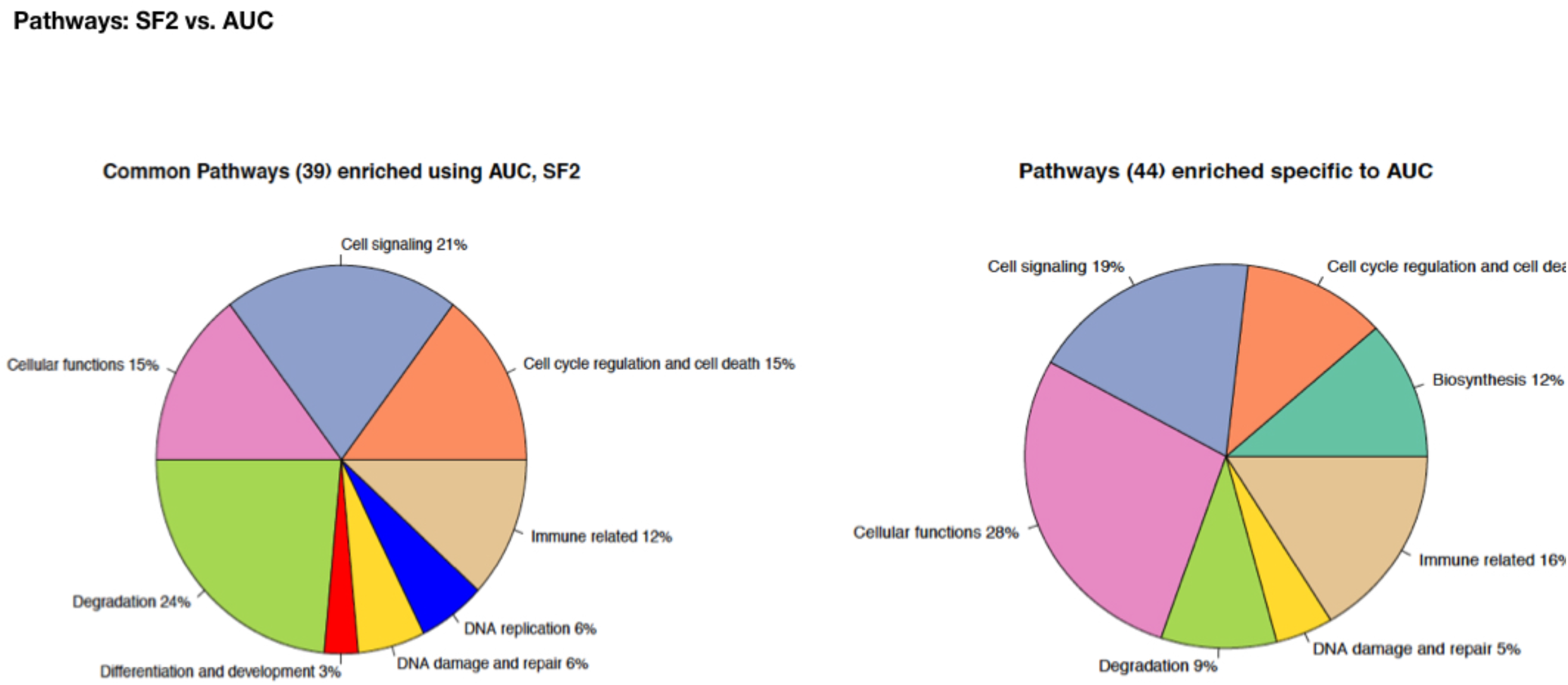
Pathway analysis comparison: SF2 vs. AUC.

**Supplementary Figure 5.**
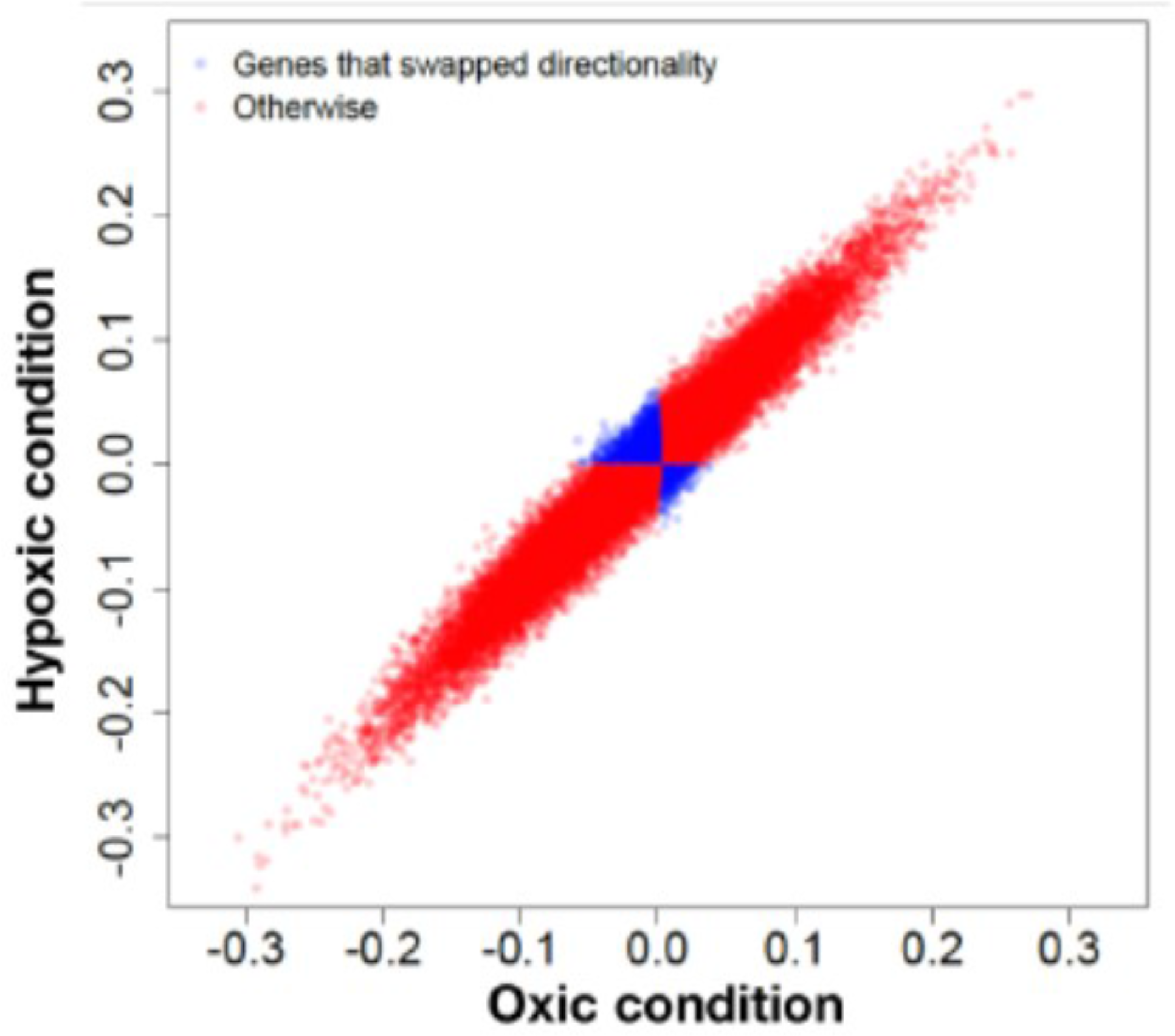
Univariate correlation between radiation response associated genes under oxic and modeled hypoxic conditions.

**Supplementary Figure 6.**
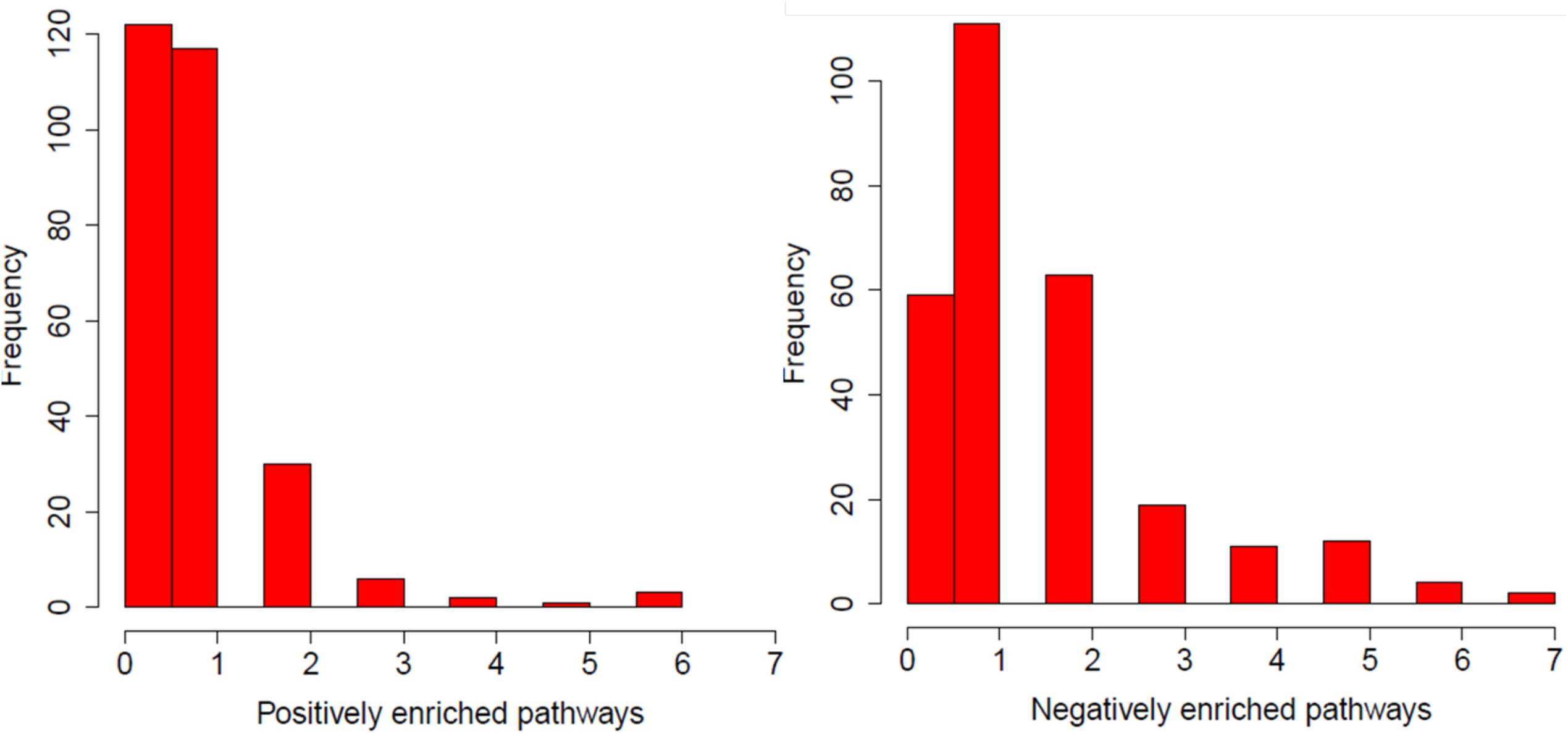
The tumour types (n=12) represented by a minimum of 15 cell lines were considered for analysis. A total of 281 pathways are enriched for FDR< 5%. Number of positively and negatively enriched pathways in each tissue.

**Supplementary Figure 7.**
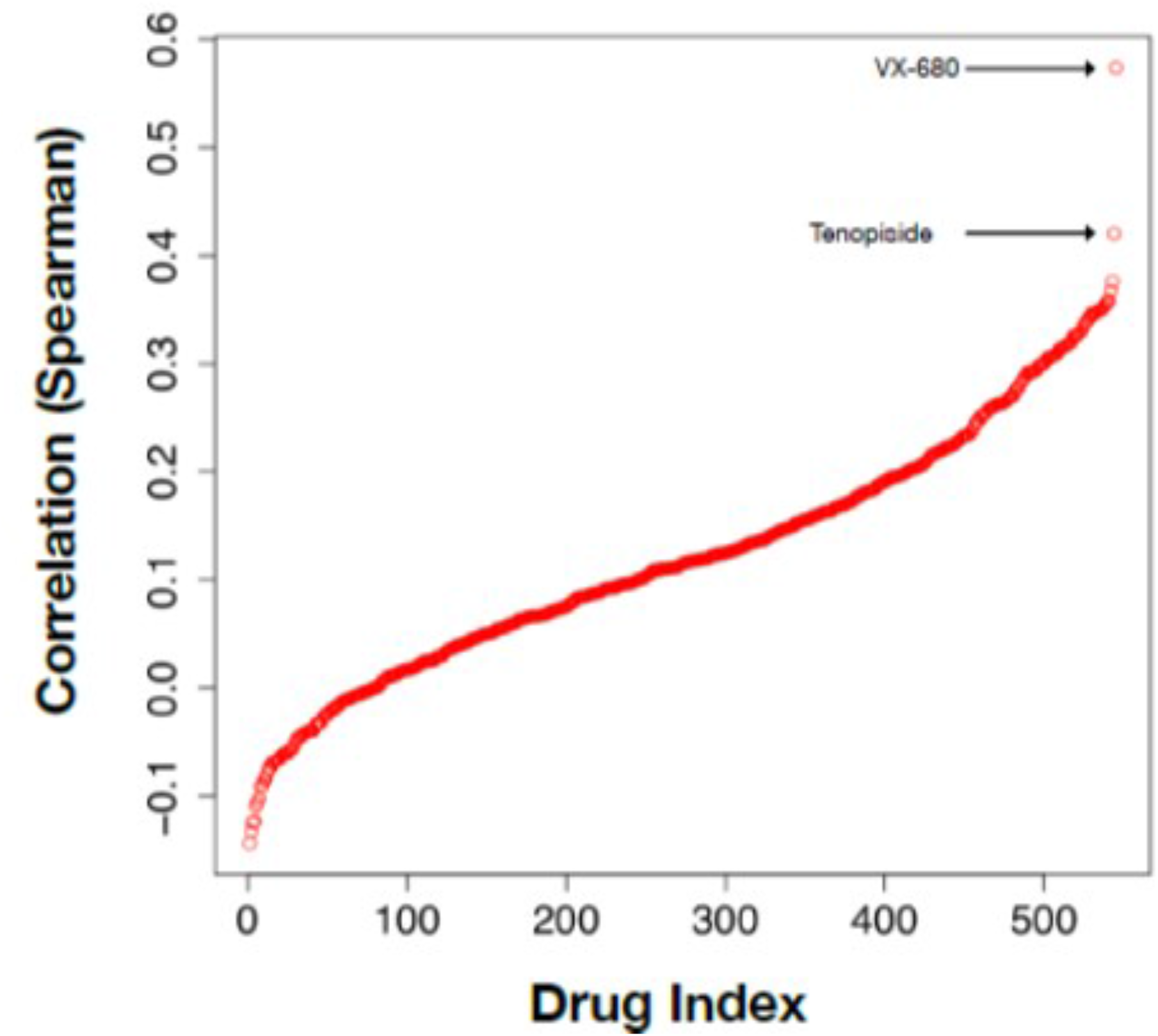
Replication of Figure 1 from Yard et al 2016, demonstrating the correlation between drug response and radiation response. Figure produced using RadioGx package.

**Supplementary Figure 8.**
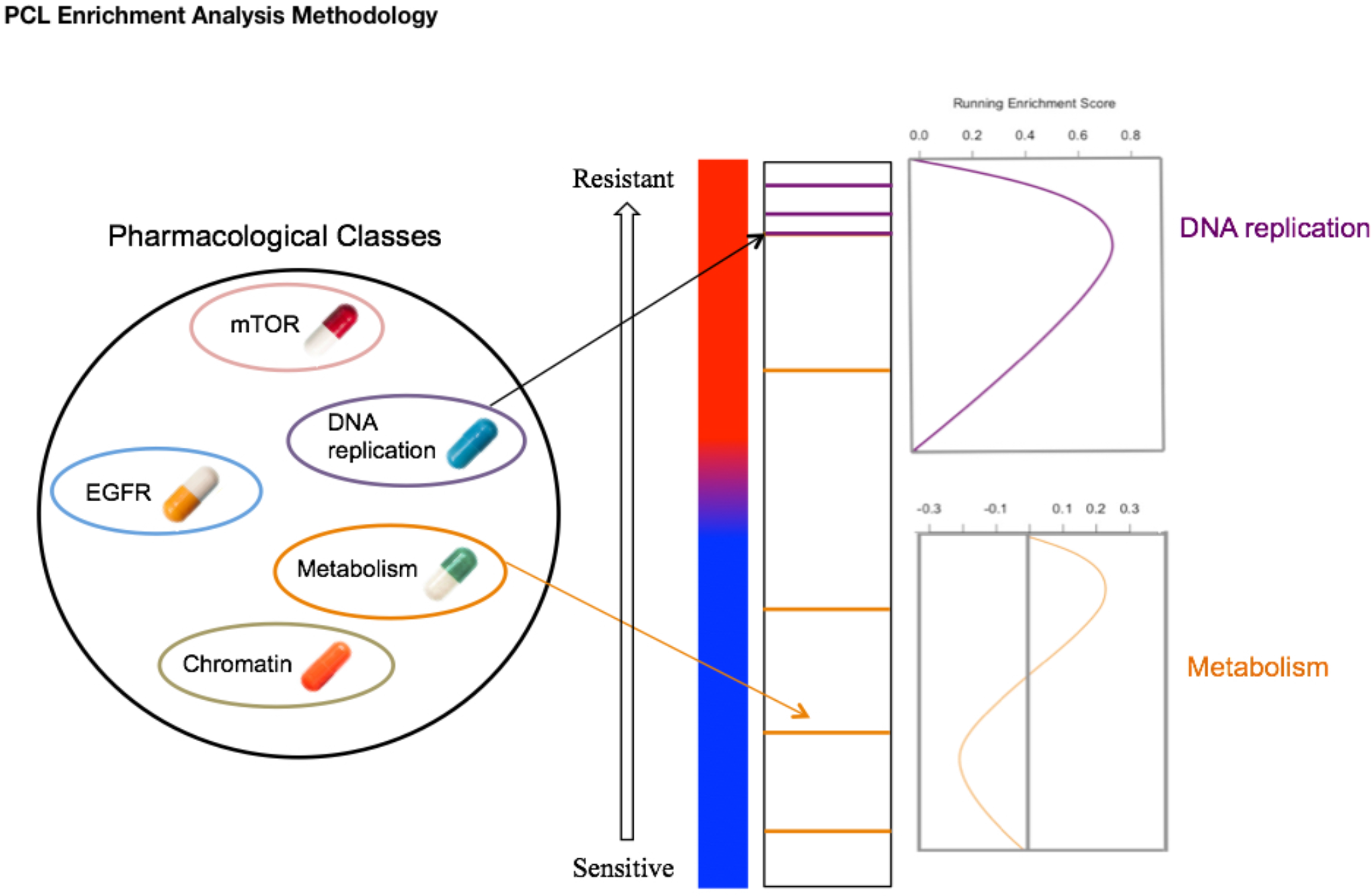
Methodology for pharmacological enrichment analysis.

